# Multidimensional neural representations of social features during movie viewing

**DOI:** 10.1101/2023.11.22.568258

**Authors:** Haemy Lee Masson, Lucy Chang, Leyla Isik

## Abstract

The social world is dynamic and contextually embedded. Yet, most studies utilize simple stimuli that do not capture the complexity of everyday social episodes. To address this, we implemented a movie viewing paradigm and investigated how the everyday social episodes are processed in the brain. Participants watched one of two movies during an MRI scan. Neural patterns from brain regions involved in social perception, mentalization, action observation, and sensory processing were extracted. Representational similarity analysis results revealed that several labeled social features (including social interaction, mentalization, the actions of others, characters talking about themselves, talking about others, and talking about objects) were represented in superior temporal gyrus (STG) and middle temporal gyrus (MTG). The mentalization feature was also represented throughout the theory of mind network, and characters talking about others engaged the temporoparietal junction (TPJ), suggesting that listeners may spontaneously infer the mental state of those being talked about. In contrast, we did not observe the action representations in frontoparietal regions of the action observation network. The current findings indicate that STG and MTG serve as central hubs for social processing, and that listening to characters talk about others elicits spontaneous mental state inference in TPJ during natural movie viewing.

## Introduction

Humans form impressions about others, such as their personality traits and social status, based on observable social cues manifesting through actions, communication, and interactions (Mehl et al., 2006; Ames et al., 2011; Quadflieg and Koldewyn, 2017). An ability to perceive a range of social cues, such as observed action (Stapel et al., 2010; Grossmann et al., 2013), social interaction (Hamlin et al., 2007; Hamlin and Wynn, 2011), speech, and communicative sounds (Dehaene-Lambertz et al., 2002; McDonald et al., 2019), emerges as early as 6 months. This early sensitivity to social stimuli observed in infancy may serve as a precursor to functional neural selectivity in adulthood (Grossmann and Johnson, 2007).

Neuroimaging studies have identified two widely recognized brain systems related to distinctive social functions: the action observation network and the mentalizing network (Van Overwalle and Baetens, 2009). Observing others’ actions activates the action observation network, including the inferior frontal gyrus (IFG), intraparietal sulcus (IPS), and superior temporal sulcus (STS) (Caspers et al., 2010; Kilner, 2011). Inferring mental states of others (i.e., theory of mind (Premack and Woodruff, 1978)) activates the mentalizing network, including the medial prefrontal cortex (mPFC), temporoparietal junction (TPJ), precuneus, and temporal pole (TP) (Gallagher and Frith, 2003; Jacoby et al., 2016). Beyond those systems, prior work has identified regions in the STS that show functional selectively to social interaction in both controlled experiments using simple stimuli (Isik et al., 2017; Walbrin et al., 2018) and in more ecologically valid studies that involve natural viewing (Lee Masson and Isik, 2021; McMahon et al., 2023). Furthermore, the STS has shown selective neural responses to visually presented social communication (McMahon et al., 2023) and speech-based social communication (Landsiedel and Koldewyn, 2023). Recent data-driven work has further suggested that communication and antisocial behavior elicit responses in the STG and MTG, respectively (Santavirta et al., 2023).

In the real social world, social communication and interaction co-occur frequently with theory of mind and action observation. However, previous research has predominantly examined these facets in isolation, resulting in gaps in our understanding of complex social processes in extended, real-world contexts. The goal of the current study is to provide an in-depth understanding of the neural mechanisms underlying complex social processes in real-world contexts by adopting a natural movie viewing approach. Specifically, by densely labeling social features of movies and employing representational similarity analysis (RSA) to the two movie fMRI datasets, we identified how social interaction, action observation, mentalization, and three types of spoken communication (characters talking about themselves, others, or things) are represented in the three neural systems implicated in social perception, action observation, and mentalization. This study builds off our prior work with the same datasets (Lee Masson and Isik, 2021) in two important ways. First by using an RSA approach versus voxelwise modeling, we can examine representational match within key hypothesized regions of interest. Second, we expanded the set of social features labeled to include richer speech labels. While different types of social interaction (e.g., helping versus hindering; arguing versus celebrating; social touch) have been investigated in cognitive neuroscience (Isik et al., 2017; Lee Masson et al., 2018; Walbrin et al., 2018; Walbrin and Koldewyn, 2019), no previous studies have explored the neural representations of different targets of speech that vary based on the spoken social content.

We find that STG and MTG are responsible for processing various social features in both movies, including all types of spoken communication (regardless of content – self, others, things), social interactions (including touch), mentalization, and others’ action. The mentalizing network, excluding the precuneus, processes the mentalization and social interaction features. Listening to characters talking about others is processed in TPJ, whereas listening to conversations revolving around objects or inanimate items is processed in TP within the mentalizing network. The frontoparietal regions of the action observation network did not represent others’ action.

## Methods

By performing RSA (Kriegeskorte et al., 2008) on two fMRI movie datasets, we evaluated how various social features are represented in the brain areas implicated in social perception, action observation, and theory of mind. To this end, using two publicly available fMRI movie datasets, where participants watched the movie Sherlock and 500 Days of Summer (Figure 1A), we analyzed neural responses of 11 brain regions to 10 different sensory and social features. Brain regions and features were selected based on prior hypotheses on social processing.

**Figure 1.**
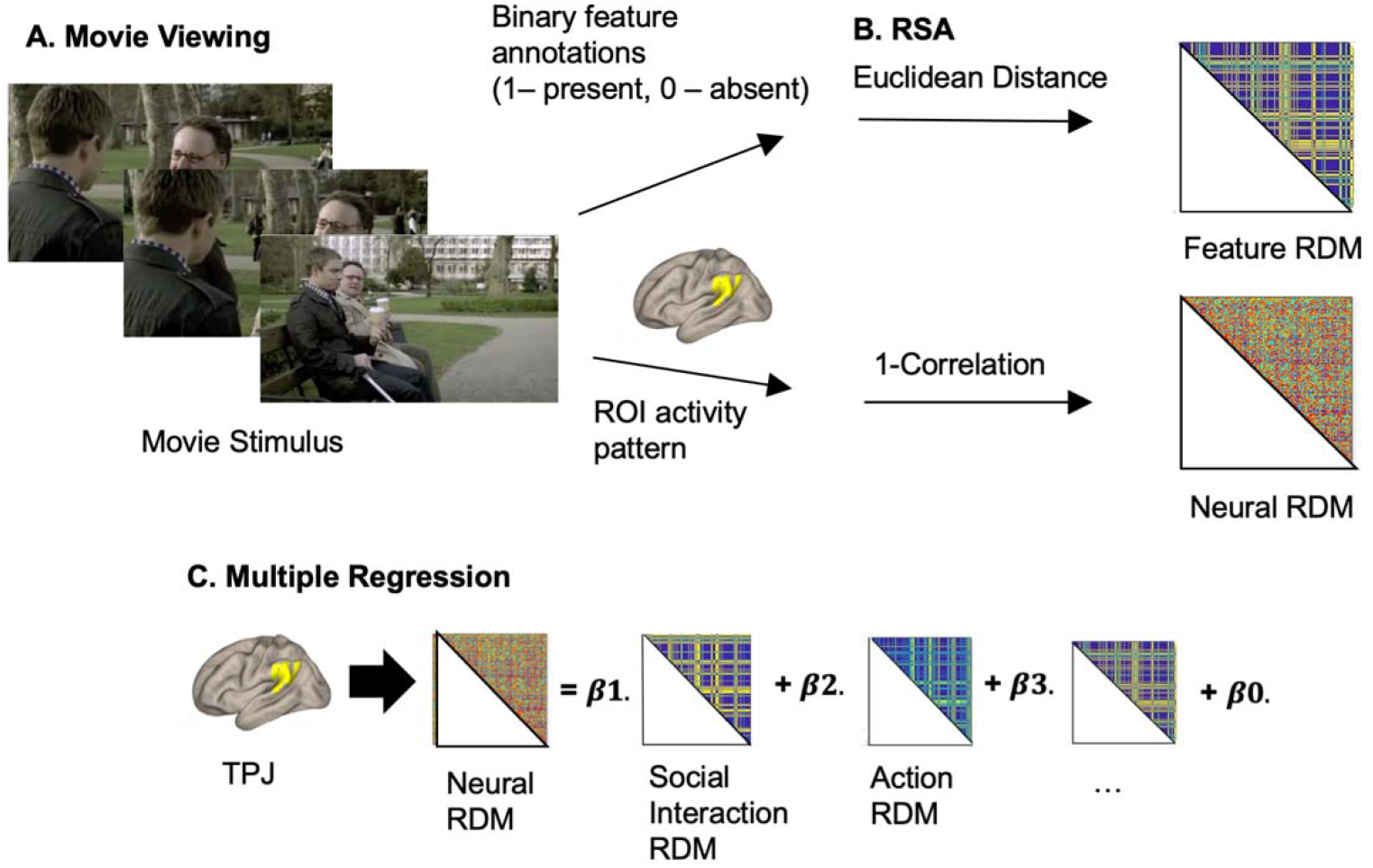
Overview of methods. A) fMRI data from participants viewing two movies–Sherlock and 500 Days of Summer– were used for analyses. Each video segment was annotated for its social content, including the presence of a social interaction, theory of mind (ToM), characters speaking about themselves, characters speaking about others, and characters speaking about things. Sensory features such as the presence of a face, audio amplitude, and visual features extracted from the fifth convolutional layer of AlexNet were also included. B) The representational dissimilarity matrices (RDMs) were created from the neural patterns for each ROI and feature. C) A multiple regression model was fit to each ROI to predict neural activity based on the different feature variables. Beta weights for each feature served as a measure of how strongly that feature explains the neural data.

### Movie feature annotations

The movies were split into 3 s segments. For the Summer movie, the opening and ending credits were cropped. This resulted in a total of 988 segments for Sherlock and 1722 segments for 500 Days of Summer. Prior to fMRI analysis, all movie segments were labeled with six social features by two raters – social interactions, mentalization, characters talking about themselves, talking about someone else, talking about something else, and actions. For the Summer movie, social touch was also labeled. The Sherlock movie rarely contained social touch scenes, so this feature was not included in Sherlock. Features were labeled 1 if the feature was present in a scene and 0 if absent. Scores was averaged across two raters.

Precisely, for the social interaction feature, we labeled scenes that involved any human-human interaction either through conversation (e.g., speaking) or action (e.g., hugging). For the spoken communication feature, we created separate vectors that represented the various talking scenes– characters talking about themselves participating in communication (lines with I/you – “You must be an army doctor”), characters talking about other characters who are not part of communication (lines with he/she/they/other character’s names – “Yeah, he’s always like that”), and characters talking about things (lines with it/object name – “On the desk there is a number.”)– in the movie. In cases where multiple communication features were present in a scene (e.g., “You got all that because you realized the case would be pink”; this scene has both characters talking about themselves and about an object – “the case”), this scene was labeled 1 for characters talking about themselves and 1 for characters talking about objects.

The presence of mentalization was labeled in each scene whenever the character was inferring another character’s emotions and thoughts (e.g., Sherlock says to Dr. Watson “Dear God, what is it like in that funny little brain of yours. It must be so boring”. In Summer, the narrator infers that “[Tom will] never be truly happy, until the day he met the one”). As labeling mentalization features based on the non-verbal expression of a character can be highly subjective, this feature is labeled solely based on the contents of the conversation. Descriptions of a character’s appearance (e.g., “She is tall and thin”) or bodily sensation (e.g., “She has been sick for 3 days”) were not labeled as mentalization.

For the action feature, only one rater labeled the actions present as there is no straightforward way to average action names across raters. The rater remained consistent with wording throughout the movie (e.g., using “speaking” instead of “talking” in all the scenes). Only the Summer movie was labeled for social touch, scenes where characters are engaging in physical contact (e.g., hugging, kissing).

To account for co-varying sensory information, we included sensory features – audio amplitude, the presence of a face, and other high-level visual features quantified as the activations extracted from the final convolutional layer of a deep neural network. We used sensory features extracted in previous studies (Aliko et al., 2020; Lee Masson and Isik, 2021). For more details see Supplementary Materials (SM).

The feature annotations were then turned into representational dissimilarity matrices (RDMs), which were used as predictors in a multiple regression model to explain the neural patterns. The feature RDM was created by calculating the pairwise Euclidean distance of each feature between all pairs of movie segments (Figure 1B). For the action RDM, identical actions were given a value of 0, while different actions (speaking vs. hugging) were given a value of 1.

### Feature correlations

To determine the correlation between each feature, we conducted a Pearson correlation analysis on all pairs of feature RDMs. We chose to use RDMs instead of raw feature annotations as this approach enabled us to establish correlations between high-dimensional visual features (256 × 13 × 13 for each scene) and other features. Results were visualized using the corrplot function in R version 3.6.3 (R Core Team, 2020). Several features were correlated across the movies. In particular, the feature capturing characters speaking about others was positively correlated with the mentalization feature in both movies. The presence of social interaction was correlated with the presence of face and action feature (Figure 2), likely reflecting the fact that ‘talking’ is the most prevalent action in both movies (Supplemental Table 1).

**Figure 2.**
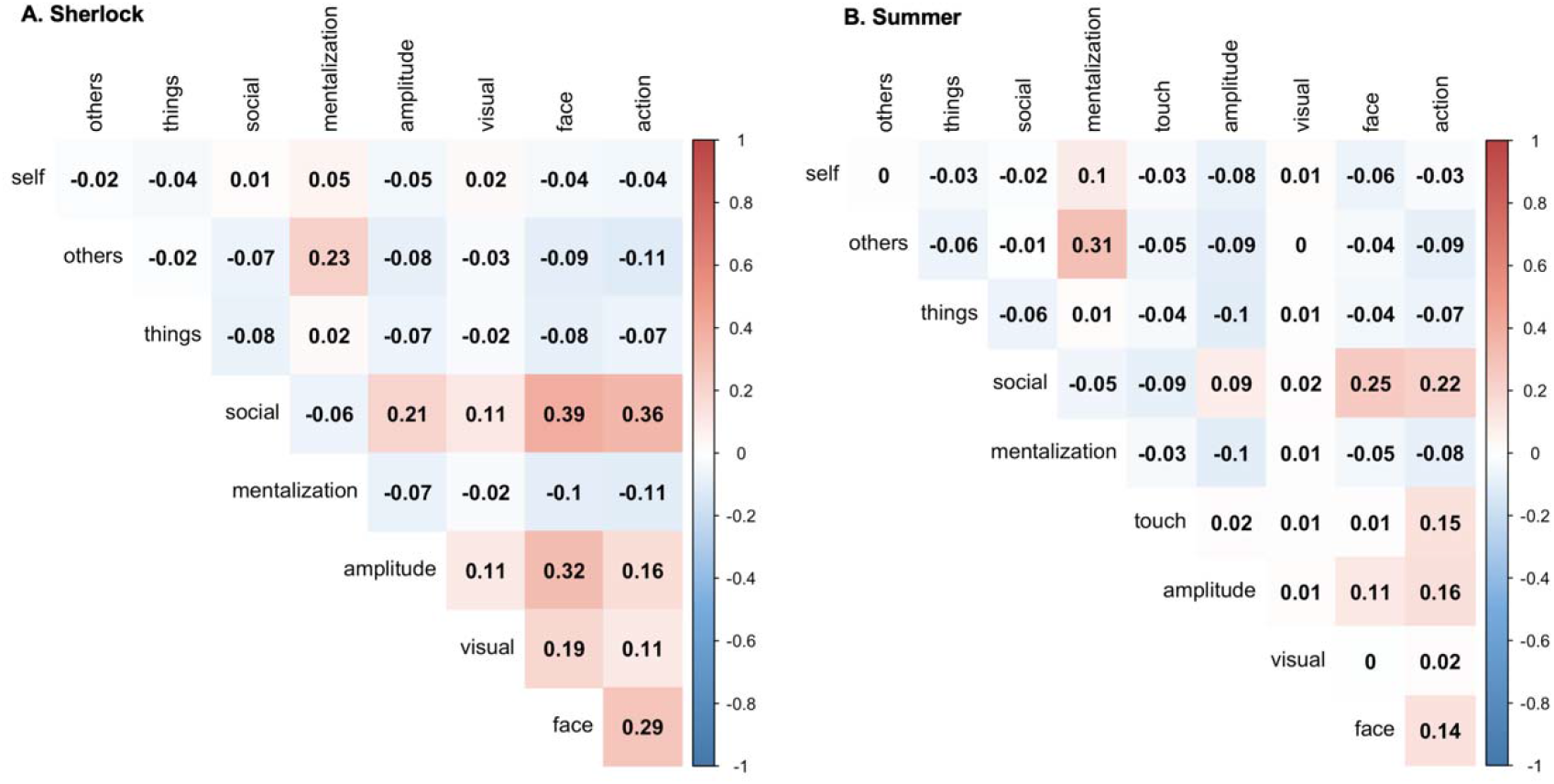
Pairwise Pearson’s correlation coefficient between feature RDMs in Sherlock (A) and Summer movies (B). Red indicates a positive correlation while the blue indicates a negative correlation. Self = characters talking about themselves, Others = characters talking about other characters, Things = characters talking about things, Social = social interaction, Visual = the fifth layer of Alexnet

### fMRI data sources

FMRI data were collected while two sets of participants watched the first episode of Sherlock BBC TV series (N = 17) and 500 Days of Summer (N = 20). Datasets are publicly available from two different studies (Chen et al., 2017; Aliko et al., 2020). The brief description of the scanning parameters can be found in SM. For the Sherlock movie, we removed one participant (Subject 5) from analysis due to missing data at the end of the scan, resulting in a total of 16 participants. For the Summer movie, we removed two participants (ID 14 and 16) as one was scanned with a different head coil and the other was given glasses only after the first run, resulting in a total of 18 participants. The studies were approved by the Princeton University Institutional Review Board and Ethics Committee of University College London, respectively. All subjects provided their written informed consent before the experiment.

### Definition of brain regions of interest (ROIs)

We conducted ROI-based analyses on fMRI data that underwent multiple preprocessing steps performed by the authors of the original study. For more details about the preprocessing steps, see SM. We measured neural representations of various social features in three well-defined networks (action observation, mentalizing, and social perception. First, anatomical ROI masks were created by using various templates. The TPJ (anterior and posterior parcels), anterior portion of mPFC (cluster 3 and 4), and posterior portion of mPFC (cluster 1 and 2) templates were taken from the connectivity-based parcellation atlas (Mars et al., 2012; Sallet et al., 2013). Following a previous study on social norm processing (Pegado et al., 2018), we separated the mPFC into two distinct ROIs. The original templates only included the right hemisphere, despite finding similar parcellation in the left hemisphere. To have bilateral ROIs, we created a mirror ROI on the left side and merged it with the original template. The STG, MTG, precuneus, opercular part of IFG, TP templates were taken from the automatic anatomical labeling atlas (Tzourio-Mazoyer et al., 2002) using PickAtlas software version 3.0.5b (Maldjian et al., 2003). The IPS mask consists of hIP1,2, and 3 templates taken from the SPM Anatomy Toolbox Version 2.2b (Eickhoff et al., 2005). We also included two control sites for visual and auditory processing. The visual brain mask consists of V1, V2, V3, V4, V5, and lateral occipital cortex extracted from the SPM Anatomy toolbox. The auditory brain mask covers primary auditory cortex extracted from the same toolbox.

Second, to select the functionally relevant voxels within each anatomical mask, we obtained brain activation maps (thresholded at z-score > 3) from Neurosynth (https://neurosynth.org/) by searching the keywords – social, action observation, and mentalizing. Using FSL from the FMRIB Software Library (Jenkinson et al., 2012), these activation maps were binarized. To define STG and MTG, we selected all voxels from the social map that were restricted to each anatomical mask (Figure 3). Anterior mPFC, posterior mPFC, TPJ, precuneus, and TP were defined by selecting all voxels from the mentalizing map within the corresponding anatomical mask. IFG and IPS were defined using the action observation map. The visual and auditory ROIs were defined with anatomical templates as they only serve as reference sites. We removed any overlapping voxels between pairs of ROIs to ensure that all ROIs were anatomically independent of each other (Table 1).

**Figure 3.**
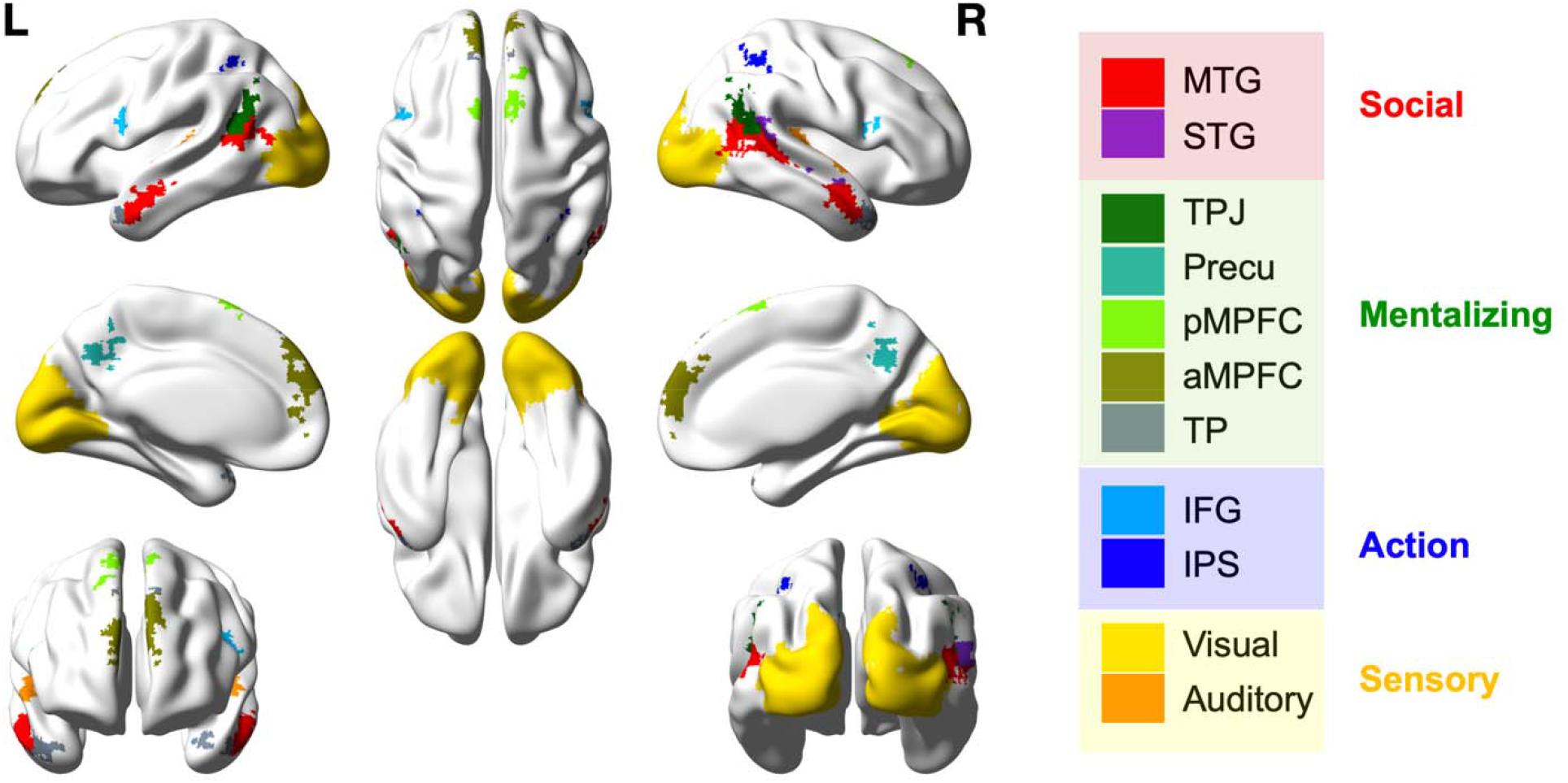
An illustration of selected ROIs visualized through BrainNetViewer (Xia et al., 2013)

**Table 1.** Overlapping voxels from ROIs pair and final ROI size.

|  | Overlapping Voxels | Final Voxel size |
| --- | --- | --- |
| STG | 11 voxels overlapping with TPJ | 77 |
| MTG | 19 with Visual, 80 with TPJ | 649 |
| TPJ | 11 with STG, 80 with MTG | 182 |
| TP | No overlap | 150 |
| Precuneus | No overlap | 221 |
| aMPFC | 8 with pMPFC | 289 |
| pMPFC | 8 with aMPFC | 98 |
| IFG | No overlap | 97 |
| IPS | No overlap | 49 |
| Auditory | No overlap | 266 |
| Visual | 19 with MTG | 4424 |

The table lists the number of overlapping voxels between ROI pairs and the final ROI size. For instance, the original STG had 11 voxels overlapping with TPJ. The last column represents the number of voxels in each ROI after the trimming.

### Neural RDMs

Prior to creating neural RDMs, BOLD signals were averaged for every two TRs (e.g., 1^st^ and 2^nd^ TR, 3^rd^ and 4^th^ TR) for Sherlock fMRI data and three TRs for Summer fMRI data to make fMRI data have a resolution of 3 s matching to feature annotations. This resulted in 988 neural patterns for Sherlock and 1722 neural pattern for Summer. To create the neural RDMs for each participant and ROI, we used the CoSMoMVPA toolbox in MATLAB (Oosterhof et al., 2016). This involved calculating the pairwise correlation distance (1 - Pearson correlation across all voxels within an ROI) of the neural patterns in response to each scene of the movie (Figure 1B). Finally, feature and neural RDMs were normalized to have the mean of 0 and a standard deviation of 1. For the subsequent analyses, the entries from the upper diagonal of RDMs were used as variables as all matrices are symmetric (Ritchie et al., 2017).

### Statistical Analysis

For the group-level statical inference, we conducted a one-tailed sign permutation test with 5000 iterations. P-values were corrected for multiple comparisons using a maximum correlation threshold across all ROIs (Nichols and Holmes, 2002).

### Inter-subject correlation as a measure of reliability

To determine reliability of the neural data, we performed a leave-one-subject-out correlation. Specifically, for each ROI, the neural RDM of one participant was correlated with the averaged neural RDMs of the other participants. A permutation test revealed that averaged correlation value across participants is above chance for all ROIs, indicating that neural data is reliable (all P_corrected values < 0.05, r-values listed in the Supplemental Table 2). Inter-subject correlation value can also be interpreted as the noise ceiling, which is expected highest correlation between neural data and other predictors, taking into account the noise present in the neural data (Nili et al., 2014).

### Multiple regression analysis

All features were assigned as predictors in the multiple regression model to explain the neural patterns in each ROI for each participant (Figure 1C). A fitlm function in MATLAB was used for this analysis. Prior to this analysis, we first checked for multicollinearity using variance inflation factors (VIF) (Marquaridt, 1970). Our analysis showed that each predictor had a VIF value of 1.3 or lower for both movies. Typically, a VIF value above 5 indicates moderate multicollinearity (Belsley, 1991). Since VIF values were well below that threshold, we assumed that critical levels of multicollinearity would not be present in our model. After the regression analysis, we performed a sign permutation test on beta values from each predictor. P values were adjusted for multiple comparisons to account for the number of ROIs tested.

### Neural pattern similarity between ROIs

Lastly, we performed a correlation analysis between the neural RDMs to identify the representational relationship between ROIs (Kriegeskorte et al., 2008). This approach differs from comparing averaged neural responses between ROIs as RSA on neural RDMs enables us to compare the representational structure across the pattern of voxels in each ROI (Pillet et al., 2020). The representational similarity between ROIs was calculated through pairwise Pearson correlation of the neural RDMs averaged across all participants for all ROI pairs. We visualized the results through multidimensional scaling (MDS) reconstruction with the mdscale function on MATLAB.

## Results

Neural patterns from 11 brain regions were extracted from two fMRI datasets recorded while subjects viewed different movies, Sherlock and 500 Days of Summer. We first computed a neural RDM for each ROI based on the pairwise similarity of each region’s response to different movie segments. To determine how brain regions involved in social perception, mentalizing, and action observation represent various social features while watching natural movies, we fit a multiple regression model to each ROI using the different feature RDMs as predictors (Figure 1). We determined that a feature was represented in the brain when it showed statistical significance in both movie fMRI datasets.

### High-level social features are represented in STG and MTG

All high-level social features – social interaction (including social touch interaction), mentalization, three types of speaking features (talking about themselves, others, and things), and action – were significantly represented in both STG and MTG (Figure 4). In contrast, visual (including the face feature) and auditory features were not consistently represented across two movies in these regions (beta-values and statistics are listed in the Supplemental Table 3).

**Figure 4.**
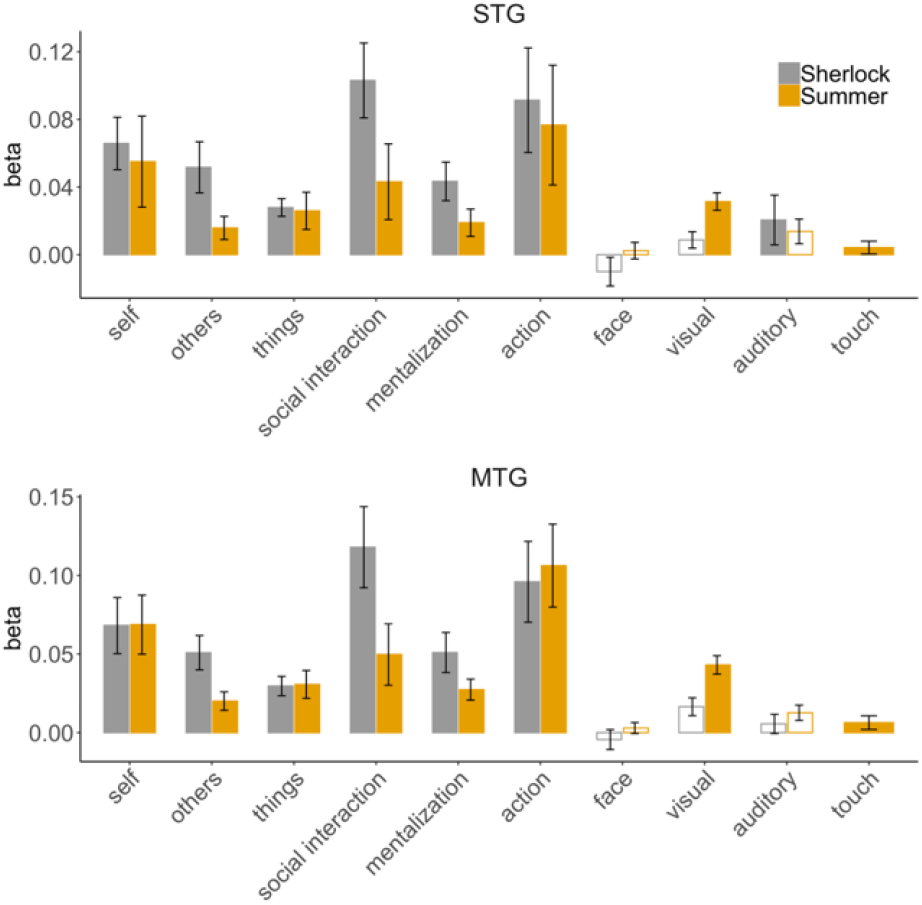
Feature representations in ROIs from the social perception network STG and MTG. The Y-axis displays the average beta value across all participants from the multiple regression model for each feature. The bar graphs include error bars to demonstrate the standard deviation. Solid bars represent statistical significance (P_corrected < 0.05), unlike empty ones.

### The mentalizing network represents the mentalization feature and characters speaking about others

As expected, we found that the mentalization feature was represented across the mentalizing network for both movies, except precuneus where the results were only significant for the Sherlock data (Figure 5). All of them represented the social interaction feature during both movies and the touch feature for Summer, with the exception of precuneus.

**Figure 5.**
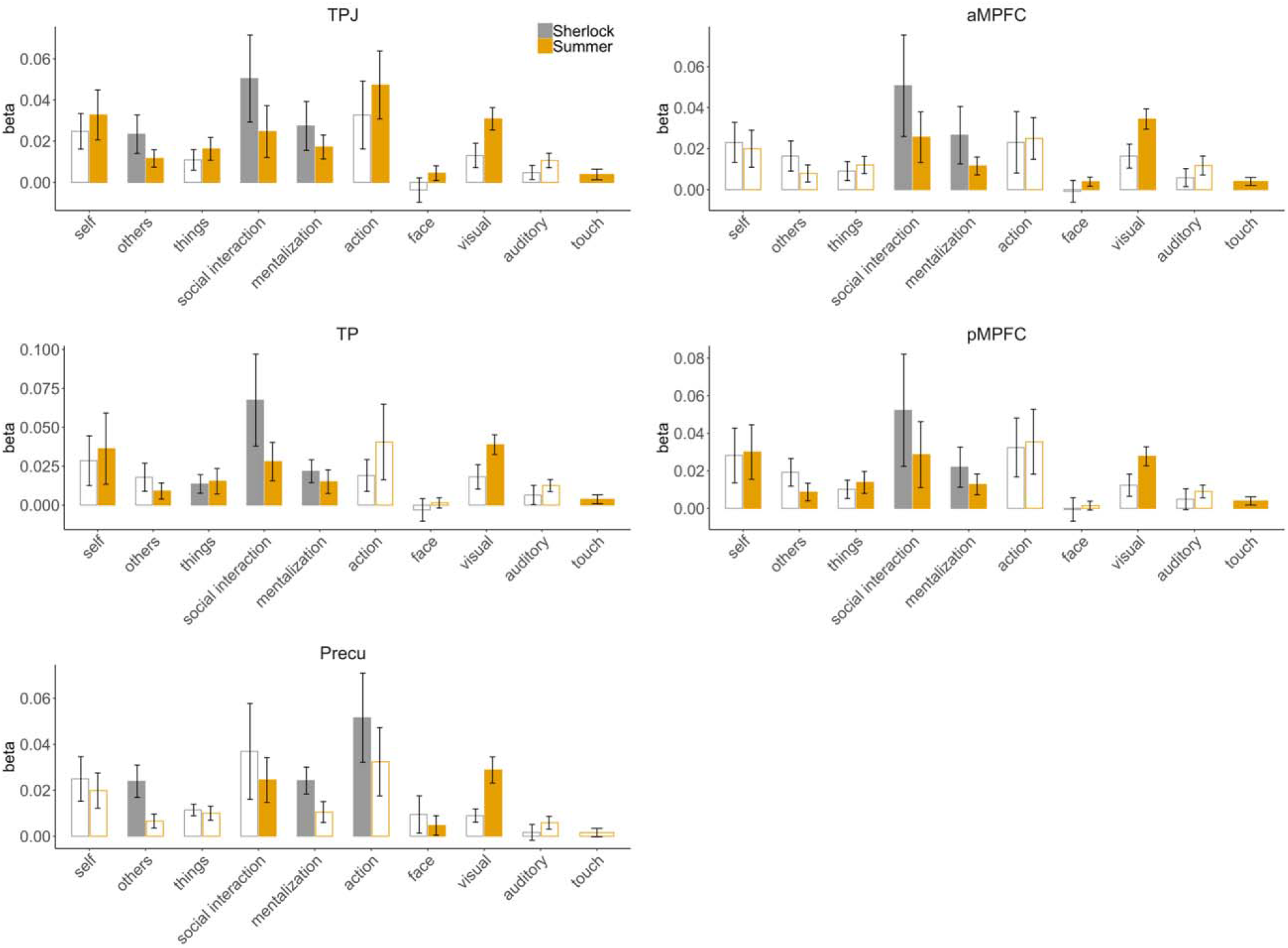
Feature representations in the mentalizing network. Plotting conventions are the same as in Figure 4.

We next examined whether the mentalizing network represented the feature capturing characters speaking about others. We hypothesized that listening to a protagonist speaking about another character would engage the mentalizing network as it may invite a viewer to make social judgements about others. However, a key difference between the mentalizing feature and this feature is that the mentalization feature only refers to the character speaking about another character’s thoughts and feeling and does not include when talking about the character’s appearance or bodily sensations, unlike the speaking about others feature which includes all of these. We found that out of all the mentalizing brain regions, only TPJ consistently represented the feature capturing speaking about others in both movies (P_corrected = 0.01 for Summer, P_corrected < 0.05 for Sherlock). Surprisingly, TP represented the feature capturing the speaking about things across both movies (P_corrected < 0.05 for both movies). No other regions in this network represented any speaking feature in a manner that was consistent across both movies.

### The action feature is not represented in the frontoparietal action observation network

Surprisingly, the beta values for observed action were not significantly above chance in either IPS or IFG (Figure 6). This is in contrast to STG and MTG which both significantly represent action information (Figure 4). The social touch feature was represented in IPS for Summer (P_corrected < 0.05). Other social features were represented in IFG (e.g., social interaction for Sherlock), but the results were not consistent across two movies.

**Figure 6.**
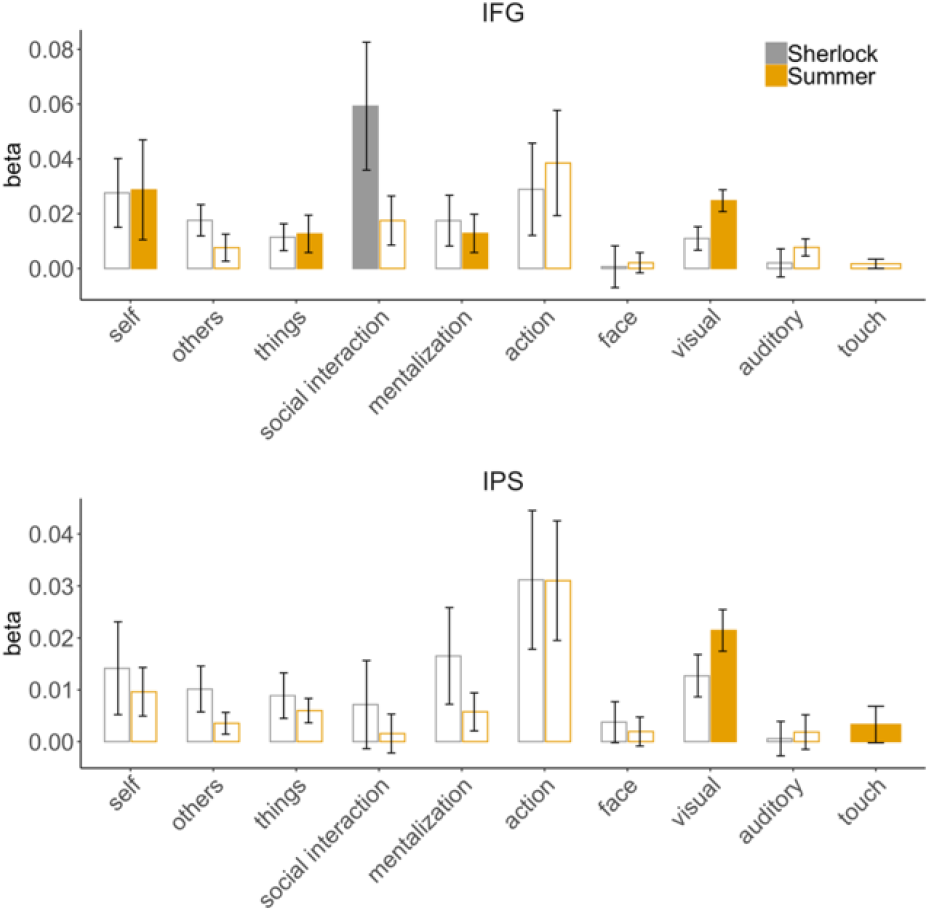
Feature representations in the action observation network. Plotting conventions are the same as in Figure 4.

**Figure 7.**
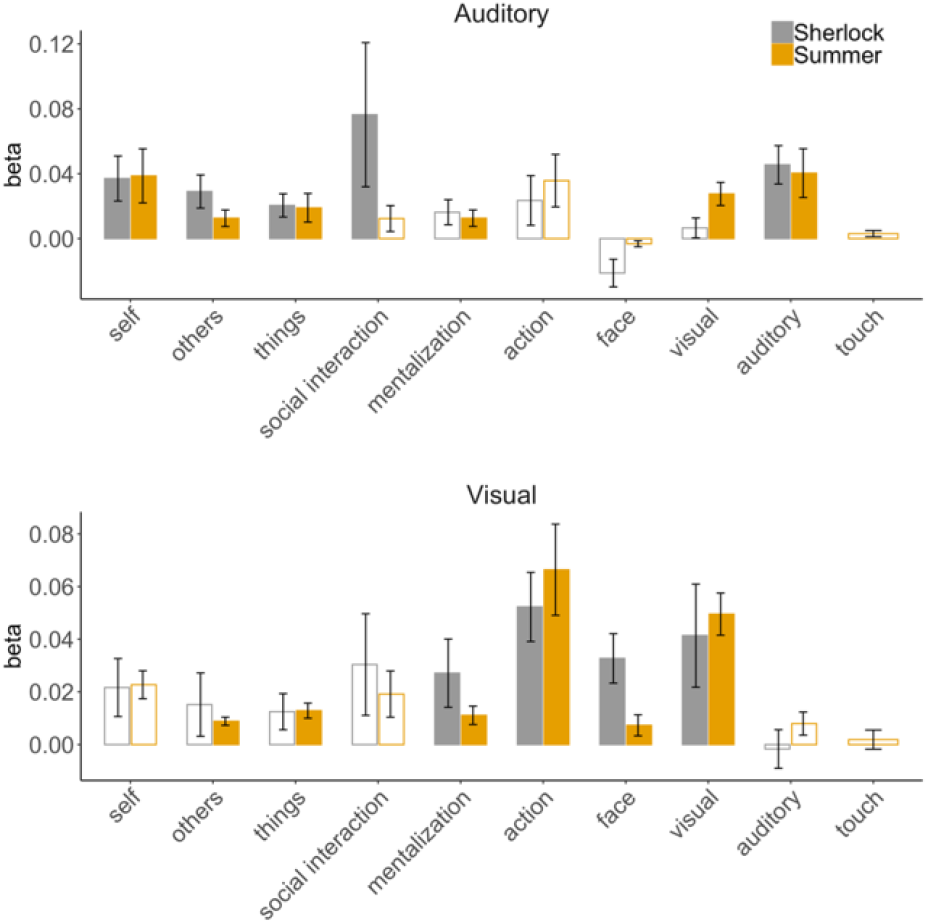
Feature representations in the visual and auditory cortex. Plotting conventions are the same as in Figure 4.

### Visual and auditory features are represented in sensory regions

As described above, we included two sensory regions – the visual cortex encompassing V1 to V5 and the auditory cortex. Mentalization, action, face, visual features extracted from the 5^th^ layer of Alexnet were represented in the visual cortex across two movies. All speaking features and amplitude of the audio were represented in the auditory cortex across two movies.

### ROIs within the same network have similar representational structure

Lastly, to understand the representational relationship between ROIs, we performed a correlation analysis on all pairs of neural RDMs. Results are visualized with MDS plots (Figure 8). We found that ROIs within each brain network, except for the action observation network, showed similar representational structure. For example, r value between STG and MTG within the social perception network was 0.83 and 0.87 for Sherlock and Summer, respectively (Supplemental Figure 1 including matrices with r values). However, IFG and IPS within the action observation network did not show strong neural pattern similarity. Instead, IFG neural patterns were more strongly correlated with those of STG (r = 0.44 and 59) and MTG (r = 0.54 and 0.65) rather than the IPS (r = 0.36 and 0.45).

**Figure 8.**
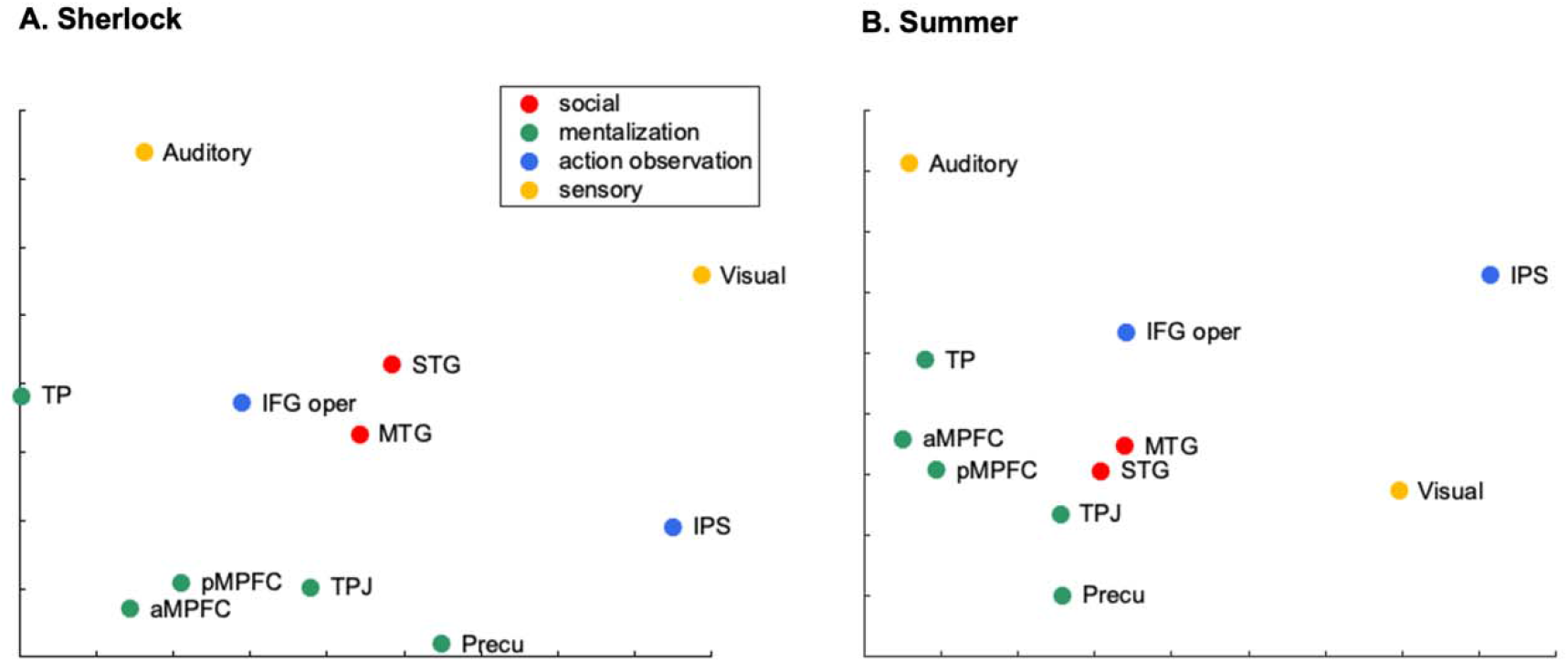
A 2D MDS plot showing the neural pattern similarity between different ROIs for Sherlock (A) and Summer (B). The distance between them is based on the 2D projection of the correlation coefficient of their neural patterns. An ROI pair with stronger correlation are closer in proximity as compared to a pair with weaker correlation. The dots on the figure are color-coded, as illustrated on the plot legend, according to which network an ROI belongs to - social perception (red), mentalization (green), action observation (blue), or sensory network (yellow).

## Discussion

We investigated the brain mechanisms underlying naturalistic social processing in a context resembling real-life situations by using publicly available two movie fMRI datasets. By densely annotating features and performing RSA (Figure 1), we identified the brain regions representing a broad range of social features (Figure 4) and those representing mentalization (Figure 5) during movie viewing. We for the first time showed how the brain represents three types of speech varying on the object being spoken about – self, others, or things (Figure 5). Lastly, comparing the representational structure of different brain regions, we identified that those in the same network had similar representations, with the exception of the action observation network (Figure 8). By analyzing two distinct sets of movie fMRI data, obtained from different participants and labs, we cross-confirmed our results, thereby enhancing the validity of the current findings.

### The STG/MTG serves as a hub for social processing during natural movie viewing

RSA and multiple regression methods revealed that the neural patterns in both the MTG and STG represented all of the selected social features in both movies (Figure 4). This finding emphasizes the critical role of the human temporal cortex in the perception of social interaction, understanding others’ actions and mental states, as well as processing different types of communication varying on the objects being spoken about. Social interaction and observed action were the most prominent social features driving the configuration of neural patterns in both regions. Additionally, these two regions share highly similar representational structures with a neural similarity of more than 0.8, though the strong correlation observed between the two regions may have been slightly overestimated due to their close anatomical proximity (Pillet et al., 2020). It is worth noting that both STG and MTG clusters in this study encompassed clusters in the STS. Despite using the anatomical template to define the STG, we found that the functionally selected voxels were mostly situated in the right STS (Supplemental Figure 2). A similar trend was observed in the MTG, except that the clusters were considerably larger and located in both hemispheres, with distinct separation between anterior and posterior regions (Supplemental Figure 3). Our findings extend previous neuroimaging studies addressing social functions within the temporal cortex. Importantly, the current study investigated novel social features that have not been previously examined, specifically, three types of communication features that vary based on the object being spoken about.

Increased STS responses have been linked to a broad range of social cognitive processes, including perceiving biological motion, goal-directed action, social interaction, and social communication, extracting meaning from speech, mental state inference, and social norm processing (Pelphrey et al., 2004; Herrington et al., 2011; Deen et al., 2015; de Heer et al., 2017; Isik et al., 2017; Pegado et al., 2018; Lee Masson and Isik, 2021; Landsiedel and Koldewyn, 2023; McMahon et al., 2023). The STG and MTG have also been implicated in social cognitive processes, including social signal detection, the integration of verbal and nonverbal social cues, extracting social-affective meaning from observed touch, extracting meaning from speech, and perceiving social communication and antisocial behavior (Price, 2012; Sugiura et al., 2014; Holler et al., 2015; Lee Masson et al., 2018; Santavirta et al., 2023). These regions show atypical neural responses to social stimuli in neurodiverse conditions, such as autism and schizophrenia (Zilbovicius et al., 2006; Redcay, 2008; Brent et al., 2014; Köchel et al., 2015; He et al., 2021). Functionally disrupting these regions results in short-term atypical social perception (Grossman et al., 2005; Akiyama et al., 2006; Saitovitch et al., 2016). While subparts of these regions have been shown to be domain-specific (Deen et al., 2015), our findings provide evidence that they respond to multiple social features. These regions may serve as hubs for social processing during natural movie viewing. Though since the selected social features are somewhat correlated (Figure 2), it might be the case that STG/MTG process their shared variance rather than each individual social feature.

### The mentalizing network represents the social interaction and mentalization feature

We observed that the neural patterns in TPJ, TP, and mPFC within the mentalizing network represented the mentalization and social interaction feature (Figure 5). Intriguingly, the social interaction feature drives the configuration of neural patterns in these regions the most strongly in both movies. Previous work with a movie viewing paradigm has also found the increased activity in the mPFC when social interaction is present in the scene (Wagner et al., 2016). Our results expand on previous work by demonstrating that not only the mPFC, but also the TPJ and TP show distinct neural patterns in response to scenes with social interaction and those without. Given that the presence of social interaction did not explain the unique variance in voxel-wise neural activity in these regions (Lee Masson and Isik, 2021), it is mostly likely that mentalization during the perception of social interaction might have influenced those findings. Previous behavioral work has suggested that when observing social interaction, individuals may naturally consider the mental states of others (Dziobek et al., 2006; Baksh et al., 2018; Grainger et al., 2019). Regarding mentalization, we replicated previous findings showing the involvement of TPJ, TP, and mPFC during the mentalization task (Dufour et al., 2013; Schurz et al., 2014; Yang et al., 2015; Jacoby et al., 2016; Moessnang et al., 2020).

Our current findings do not entirely confirm our previous work on the same dataset, employing a different methodology, where we predicted the magnitude of voxel-wise neural responses (Lee Masson and Isik, 2021). In contrast to our prior work showing unique selectivity to social interaction in the precuneus across two movies, we did not observe its neural patterns representing social interaction in the Sherlock movie. This slightly discrepancy may be due to the relatively small cluster identified in our previous work (the number of voxels = 25 in the Sherlock fMRI) compared to the precuneus region defined in the present study (the number of voxels = 221) or discrepancies between univariate and those from multivoxel pattern analyses (Pillet et al., 2020). This feature may solely account for the voxel-wise activity without explaining the underlying patterns of those voxels.

Lastly, while we replicated our findings regarding social interaction and mentalization features across two movies, we noticed a discrepancy in the remaining features. Specifically, we observed more social features that were significantly represented in these regions in the Summer fMRI data. Due to the vast differences between the two movies in terms of genre (crime versus romance), duration (40 versus 120 minutes), and the relationship between characters (colleagues versus romantic partners), it is challenging to pinpoint the exact factors that contributed to these discrepancies.

### Processing different types of spoken communication

The current findings on the representations of all three types of speech in STG, MTG, and the auditory cortex align well with the previous literature. The temporal cortex has been long implicated in speech comprehension (Crinion et al., 2003; Lindenberg and Scheef, 2007; Leonard and Chang, 2014) and listening to dialogs between people activates those areas (Landsiedel and Koldewyn, 2023; Santavirta et al., 2023). Intriguingly, we observed subtle differences in how each speaking feature was represented in the mentalization network. The neural patterns in TPJ exhibited sensitivity to the information about others, whereas the neural patterns in TP showed sensitivity to the information about things. This finding suggests a differentiation in the neural processing of social versus non-social information when listening to others’ conversations.

Our finding shows that the TPJ is not only activated when we directly engage in inferring others’ mental states during social interaction, but also when we passively listen to someone else talking about others during movie viewing. This suggests that when someone talks about another person, a listener may spontaneously make inferences about the person being discussed, even when the spoken content is not about mental states. In contrast, the anterior temporal lobe, which includes TP, plays a key role in semantic processing (Patterson et al., 2007; Gesierich et al., 2012; Visser et al., 2012). These studies have predominantly investigated the ATL responses to controlled visual stimuli. Our result extends the previous findings by demonstrating the neural representations of things in TP when listening to other people’s natural conversations.

### Action observation network

The frontoparietal regions in the action observation network do not appear to play much role in representing social features in a manner that generalized across movies. Moreover, even the action feature was not consistently represented in this network. Prior work with controlled stimuli has suggested that representations in these regions may not generalize across scenes with different kinematic patterns or with variable visual information (Wurm and Lingnau, 2015), which may explain their lack of consistent response in natural movies that involve highly variable visual information. The lack of action representation within the action network may also be linked to the high number of speaking scenes present in the movie, which makes the action category rather simplistic, reducing it to essentially speaking vs. other action categories. Future research may use movies with fewer speaking scenes and more action variability to verify whether fine-detailed action categories are represented in this network during natural movie viewing.

## Conclusion

Our study investigated the neural representations of various social features in natural movies, a setting closer to real-life social scenarios than typical experiments, including social interaction perception, listening to others’ conversations, action observation, and mentalization. Our findings highlight the temporal cortex as a hub for naturalistic social processing and suggest that different cognitive processes may come into play depending on whether a conversation concerns the speaker themselves, others, or inanimate objects. Moreover, we found high similarities in the activity patterns of brain regions responsible for social perception and mentalization, suggesting a collaborative effort among these regions in natural settings to combine various social cues Our study draws generalized conclusions from two distinct fMRI datasets, improving the reliability of the current findings. Future research may explore the temporal dimension of these social processes, which would shed light on the order of computational steps unfolding within these brain networks.

## Conflict of interest

The authors declare no competing financial interests.

## Acknowledgements

This work was supported with funds from The Clare Boothe Luce Program for Women in STEM.

## Data and code availability statement

fMRI data is available from the original authors for Sherlock (https://dataspace.princeton.edu/handle/88435/dsp01nz8062179) and Summer (https://openneuro.org/datasets/ds002837/, https://www.naturalistic-neuroimaging-database.org/index.html). Annotations are available at https://osf.io/98rfv/. All code for RSA is available at https://github.com/lchang31/MultipleRegression.

## Notes

### Competing Interest Statement

The authors have declared no competing interest.

