## Supplemental Materials for "Multidimensional neural representations of social features during movie viewing"

1. **Movie annotations of sensory features**

For the Sherlock movie, those features were extracted at a resolution of 1.5 s, which we downsampled to 3 s to match with a resolution of how social features were labeled. To briefly describe, the visual features were extracted with the Pytorch (version 1.4.0) implementation of AlexNet (Krizhevsky et al., 2012), pretrained on the ImageNet dataset (Russakovsky et al., 2015). Visual features of each scene were captured with the activations of all units from the fifth layer right before the rectified linear activation function and the max-pooling layer during the forward pass. The fifth layer captures low, mid, and high-level visual information shown in the movies (Lee Masson and Isik, 2021). The output size of the fifth layer was 256 × 13 × 13 for each scene and vectorized for subsequent RSA analyses. The presence of face in a scene was annotated using the Amazon Rekognition service (<https://aws.amazon.com/rekognition/>). Lastly, the audio amplitude was extracted with the audioread built-in function in MATLAB (R2020a, The Mathworks, Natick, MA). The values over the audio samples and channels for each movie segment were averaged.

2. **fMRI data preprocessing**

In the first study, “Sherlock”, whole-brain images (27 slices, a voxel size of 4 × 3 × 3 mm3) were obtained on a 3T Siemens Skyra scanner with a 20-channel head coil and an echo-planar (EPI) T2∗-weighted sequence. The acquisition parameters were as follows: repetition time (TR) = 1500 ms, echo time (TE) = 28 ms, flip angle (FA) = 64°, and field of view (FOV) = 192 × 192 mm2. In the second study, “Summer”, whole-brain images (40 slices, a voxel size of 3.2 × 3.2 × 3.2 mm3) were obtained on a 1.5T Siemens MAGNETOM Avanto with a 32-channel head coil and a multiband EPI sequence. The acquisition parameters were as follows: multiband factor = 4, no in-plane acceleration, TR = 1000 ms, TE = 54.8 ms, and FA = 75°.

The Sherlock fMRI data underwent several preprocessing steps, including correction for slice timing and motion, linear detrending, temporal high-pass filtering (140 s cut off), spatial normalization to a Montreal Neurological Institute (MNI) space with a re-sampling size of 3 × 3 × 3 mm^3^, and spatial smoothing with a 6-mm full width at half maximum (FWHM) Gaussian kernel. fMRI data were shifted by 4.5 s (3 TRs) from the stimulus onset to account for the delay in hemodynamic response. Lastly, the blood oxygenation level dependent (BOLD) signals were standardized through z-scoring.

The Summer fMRI data underwent preprocessing procedures identical to those employed in the Sherlock study. Furthermore, the authors introduced additional steps, including despiking and rescaling the timeseries BOLD signals to a normalized range of 0 to 1. The BOLD signals were also detrended based on run lengths, head-motion parameters, and averaged BOLD signals in white matter and cerebrospinal fluid regions. Lastly, timing correction was applied to ensure that the fMRI timeseries and the movie were aligned. To this end, we shifted Summer fMRI data by 4 s (4 TRs) from the stimulus onset to account for the delay in hemodynamic response.

**Table 1** Each movie’s top 20 action labels and how frequently they occur in each movie.

| Sherlock Action | Sherlock Counts | Summer Action | Summer counts |
| --- | --- | --- | --- |
| talking | 388 | talking | 604 |
| walking | 73 | walking | 182 |
| no action | 55 | no action | 58 |
| sitting, talking | 19 | looking at someone | 39 |
| walking, talking | 16 | looking at each other | 32 |
| telephoning | 15 | pointing | 31 |
| sitting | 12 | sitting | 30 |
| playing music | 8 | singing | 30 |
| lying down | 8 | kissing | 28 |
| driving | 8 | laughing | 27 |
| analyzing | 8 | dancing | 26 |
| staring | 7 | nodding | 23 |
| reading message | 7 | drawing | 23 |
| turning around | 6 | turning head | 21 |
| observing | 6 | drinking | 21 |
| crying | 5 | lying | 21 |
| giving report | 5 | smiling | 19 |
| talking, walking | 5 | running | 18 |
| standing | 5 | giving | 18 |
| looking down | 4 | driving | 17 |

Full action labels can be found in <https://osf.io/98rfv/>.

**Table 2** The reliability measured with leave one subject out correlation

|  | STG | MTG | TPJ | TP | Precu | aMPFC | pMPFC | IFG | IPS | Auditory | Visual |
| --- | --- | --- | --- | --- | --- | --- | --- | --- | --- | --- | --- |
| Sherlock | 0.52*** | 0.60*** | 0.38** | 0.34* | 0.41** | 0.33* | 0.33* | 0.36* | 0.28* | 0.35* | 0.53*** |
| Summer | 0.42*** | 0.56*** | 0.40*** | 0.30* | 0.40*** | 0.33** | 0.31* | 0.30* | 0.33** | 0.36** | 0.53*** |

For each ROI, the neural RDM of one participant was correlated with the averaged neural RDMs of the other participants. Correlation r-values were averaged across participants. *** P_corrected < 0.001, ** P_corrected < 0.01, * P_corrected < 0.05

**Table 3a** Beta values from the multiple regression model for the Sherlock movie

|  | Self | Others | Things | Social Interaction | Mentalization | Auditory | Visual | Face | Action |
| --- | --- | --- | --- | --- | --- | --- | --- | --- | --- |
| STG | 0.07^***^ | 0.05^***^ | 0.03^***^ | 0.1^***^ | 0.04^***^ | 0.02^*^ | 0.01 | -0.01 | 0.09^***^ |
| MTG | 0.07^***^ | 0.05^***^ | 0.03^***^ | 0.12^***^ | 0.05^***^ | 0.01 | 0.02 | 0 | 0.1^***^ |
| TPJ | 0.02 | 0.02^*^ | 0.01 | 0.05^*^ | 0.03^*^ | 0 | 0.01 | 0 | 0.03 |
| TP | 0.03 | 0.02 | 0.01^*^ | 0.07^**^ | 0.02^*^ | 0.01 | 0.02 | 0 | 0.02 |
| Precu | 0.02 | 0.02^*^ | 0.01 | 0.04 | 0.02^*^ | 0 | 0.01 | 0.01 | 0.05^*^ |
| aMPFC | 0.02 | 0.02 | 0.01 | 0.05^*^ | 0.03^*^ | 0.01 | 0.02 | 0 | 0.02 |
| pMPFC | 0.03 | 0.02 | 0.01 | 0.05^*^ | 0.02^*^ | 0 | 0.01 | 0 | 0.03 |
| IFG | 0.03 | 0.02 | 0.01 | 0.06^*^ | 0.02 | 0 | 0.01 | 0 | 0.03 |
| IPS | 0.01 | 0.01 | 0.01 | 0.01 | 0.02 | 0 | 0.01 | 0 | 0.03 |
| Auditory | 0.04^*^ | 0.03^**^ | 0.02^**^ | 0.08^**^ | 0.02 | 0.05^***^ | 0.01 | -0.02 | 0.02 |
| Visual | 0.02 | 0.02 | 0.01 | 0.03 | 0.03^*^ | 0 | 0.04^***^ | 0.03^***^ | 0.05^*^ |

Asterisks denote corrected P values from permutation test (* P < 0.05, ** P < 0.01. *** P < 0.001).

**Table 3b** Beta values from the multiple regression model for the 500 Days of Summer movie

|  | Self | Others | Things | Social Interaction | Mentalization | Touch | Visual | Auditory | Face | Action |
| --- | --- | --- | --- | --- | --- | --- | --- | --- | --- | --- |
| STG | 0.06^***^ | 0.02^***^ | 0.03^***^ | 0.04^***^ | 0.02^**^ | 0.004^*^ | 0.03^**^ | 0.01 | 0 | 0.08^***^ |
| MTG | 0.07^***^ | 0.02^***^ | 0.03^***^ | 0.05^***^ | 0.03^***^ | 0.01^***^ | 0.04^***^ | 0.01 | 0 | 0.11^***^ |
| TPJ | 0.03^*^ | 0.01^**^ | 0.02^*^ | 0.02^*^ | 0.02^**^ | 0.004^*^ | 0.03^**^ | 0.01 | 0.004^*^ | 0.05^*^ |
| TP | 0.04^*^ | 0.01^*^ | 0.02^*^ | 0.03^*^ | 0.01^*^ | 0.004^*^ | 0.04^**^ | 0.01 | 0 | 0.04 |
| Precu | 0.02 | 0.01 | 0.01 | 0.02^*^ | 0.01 | 0 | 0.03^**^ | 0.01 | 0.005^**^ | 0.03 |
| aMPFC | 0.02 | 0.01 | 0.01 | 0.03^*^ | 0.01^*^ | 0.004^*^ | 0.03^**^ | 0.01 | 0.004^*^ | 0.02^*^ |
| pMPFC | 0.03^*^ | 0.01^*^ | 0.01^*^ | 0.03^*^ | 0.01^*^ | 0.004^*^ | 0.03^**^ | 0.01 | 0 | 0.04 |
| IFG | 0.03^*^ | 0.01 | 0.01^*^ | 0.02 | 0.01^*^ | 0 | 0.02^*^ | 0.01 | 0 | 0.04 |
| IPS | 0.01 | 0 | 0.01 | 0 | 0.01 | 0.003^*^ | 0.02^*^ | 0 | 0 | 0.03 |
| Auditory | 0.04^*^ | 0.01^**^ | 0.02^**^ | 0.01 | 0.01^*^ | 0 | 0.03^**^ | 0.04^***^ | 0 | 0.04 |
| Visual | 0.02 | 0.01^*^ | 0.01^*^ | 0.02 | 0.01^*^ | 0 | 0.05^***^ | 0.01 | 0.01^***^ | 0.07^**^ |

Asterisks denote corrected P values from permutation test (* P < 0.05, ** P < 0.01. *** P < 0.001).

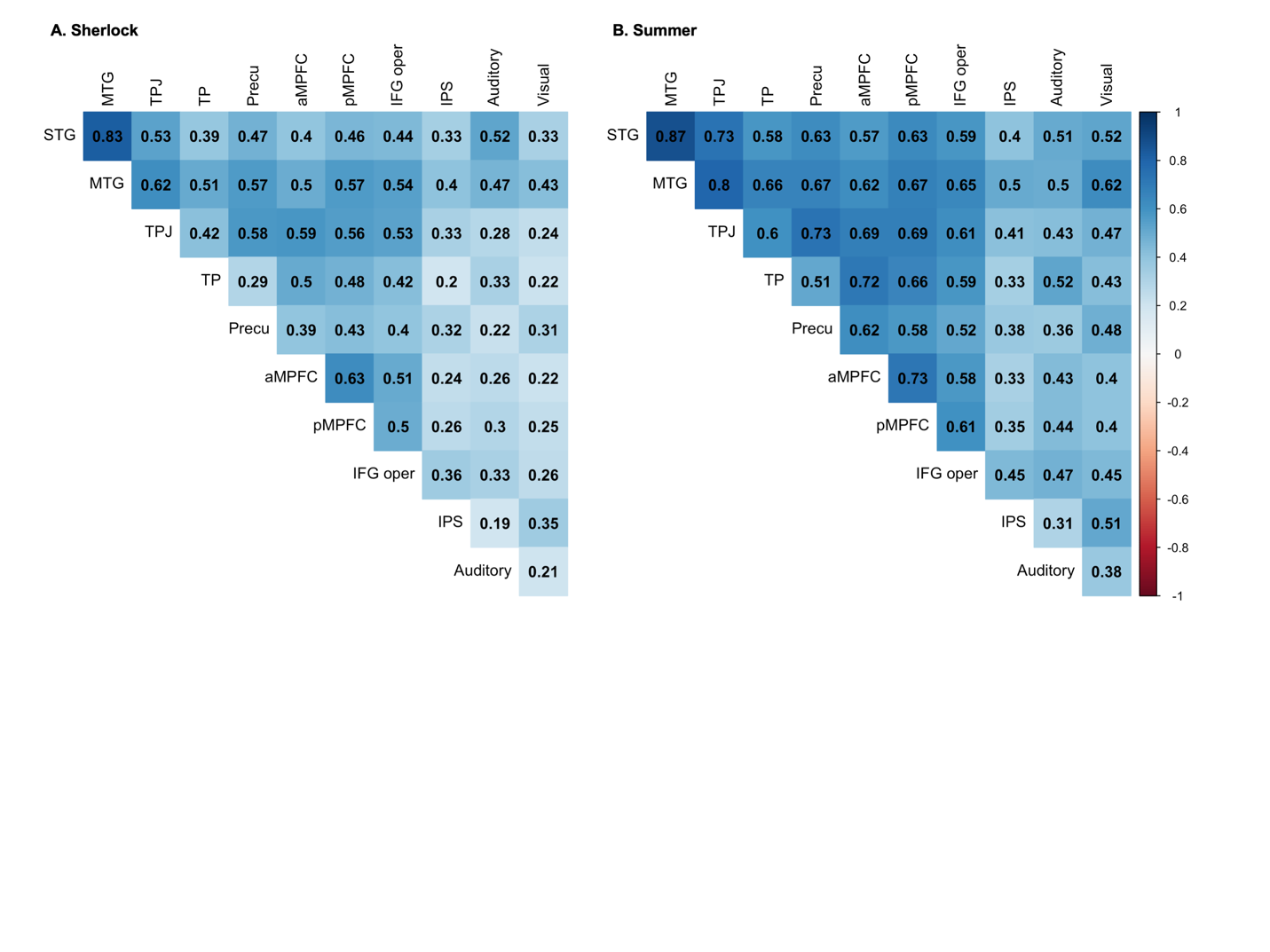

**Figure 1** Pairwise Pearson’s correlation coefficient between neural RDMs from ROIs in Sherlock (A) and Summer movies (B). Blue indicates a positive correlation while the red indicates a negative correlation.

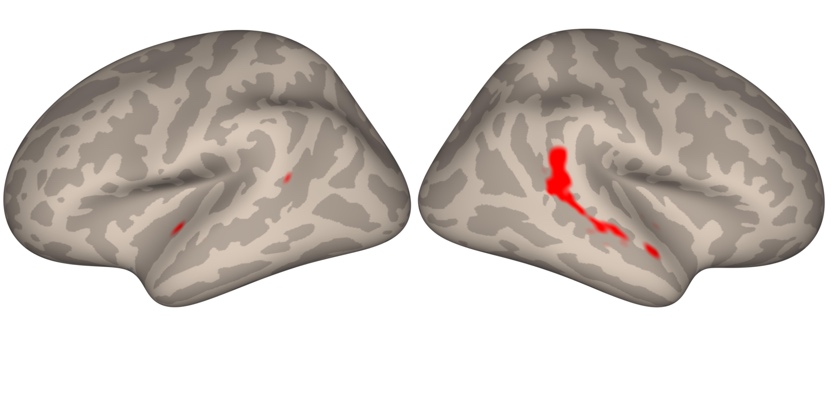

**Figure 2** The STG mask is visualized on the cortical surface in red.

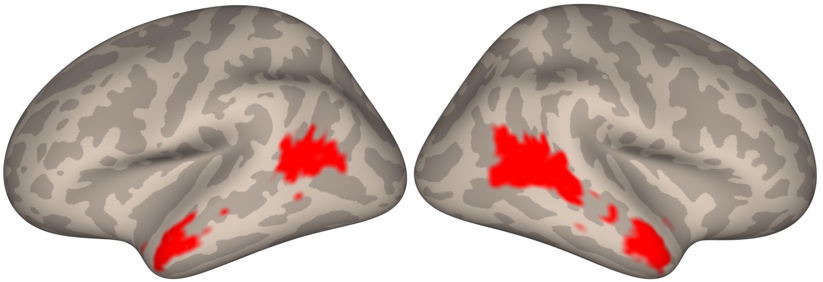

**Figure 3** The MTG mask is visualized on the cortical surface in red.
